# Genomic and machine learning-based screening of aquaculture associated introgression into at-risk wild North American Atlantic salmon (*Salmo salar*) populations

**DOI:** 10.1101/2022.11.23.517511

**Authors:** Cameron M. Nugent, Tony Kess, Matthew K. Brachmann, Barbara L. Langille, Melissa K. Holborn, Samantha V. Beck, Nicole Smith, Steven J. Duffy, Sarah J. Lehnert, Brendan F. Wringe, Paul Bentzen, Ian R. Bradbury

## Abstract

The negative genetic impacts of gene flow from domestic to wild populations can be dependent on the degree of domestication and exacerbated by the magnitude of pre-existing genetic differences between wild populations and the domestication source. Recent evidence of European ancestry within North American aquaculture Atlantic salmon (*Salmo salar*) has elevated the potential impact of escaped farmed salmon on often at-risk wild North American salmon populations. Here we compare the ability of single nucleotide polymorphism (SNP) and microsatellite (SSR) marker panels of different sizes (7-SSR, 100-SSR, and 220K-SNP) to detect introgression of European genetic information into North American wild and aquaculture populations. Linear regression comparing admixture predictions for a set of individuals common to the three data sets showed that the 100-SSR panel and 7-SSR panels replicated the full 220K-SNP-based admixture estimates with low accuracy (r^2^ of 0.64 and 0.49 respectively). Additional tests explored the effects of individual sample size and marker number, which revealed that ~300 randomly selected SNPs could replicate the 220K-SNP admixture predictions with greater than 95% fidelity. We designed a custom SNP panel (301-SNP) for European admixture detection in future monitoring work and then developed and tested a Python package, SalmonEuAdmix (https://github.com/CNuge/SalmonEuAdmix), that uses a deep neural network to make *de novo* estimates of individuals’ European admixture proportion without the need to conduct complete admixture analysis utilizing baseline samples. The results demonstrate the mobilization of targeted SNP panels and machine learning in support of at-risk species conservation and management.

## Introduction

Losses of biodiversity and accelerating rates of species extinction have now been documented across the globe (Barnosky *et al*. 2011). Attempts to stem this tide of inter- and intraspecific loss requires a robust understanding of causal factors involved, which is often lacking. Wild populations of Atlantic salmon (*Salmo salar*) in Atlantic Canada are highly valued for their ecological, cultural, and commercial importance (DFO 2019). Across the North Atlantic, more than 60% of salmon populations show evidence of decline in recent decades (Lehnert *et al*. 2019). Within Canadian waters, large population declines have been observed, with abundance estimated to have fallen by 50% in the last half century; the largest declines have been seen in populations of the Bay of Fundy, Southern Nova Scotia, and Southern Newfoundland (COSEWIC 2010; DFO 2019). The causes of decline are largely unknown, but possibilities include climate change (*e.g*., Nicola *et al*. 2018; Lehnert *et al*. 2019), fishery exploitation (*e.g*., Bradbury *et al*. 2015; Dadswell *et al*. 2021), predation (*e.g*., Daniels *et al*. 2018; Strøm *et al*. 2019), and interactions with salmon aquaculture (*e.g*., Glover *et al*. 2017; Wringe *et al*. 2018; Bradbury *et al*.2020). Ultimately, the resolution of these causal factors will be key to the prevention of further extirpation and the success of any recovery or restoration efforts.

Interbreeding of Atlantic salmon aquaculture escapees with wild salmon has been identified as a significant threat to the species’ persistence and stability in the wild (Forseth *et al*. 2017). Both hybridization and subsequent introgression have been observed in wild populations across the North Atlantic (*e.g*., Karlsson *et al*. 2016; Wringe *et al*. 2018; Gilbey *et al*. 2021) and have been shown to reduce population viability through maladaptive genetic changes to wild stocks (Sylvester *et al*. 2019; Bolstad *et al*. 2017, 2021). Evidence of profound genomic differences (*e.g*., Lehnert *et al*. 2019, 2020) as well as behavioral, and physiological differences (*e.g*., Islam *et al*. 2021) between European and North American salmon support the hypothesis that the negative effects of European escapees in North America likely exceed those of North American individuals (Bradbury *et al*. 2022). As a result, restrictions on the use of European salmon in North America have been in place since the late 1990s (Baum *et al*. 1998; Porter *et al*. 1998; DFO 2016). Nonetheless, mounting evidence suggests the continued presence of Atlantic salmon with European ancestry in: North American aquaculture salmon, escapees, and wild salmon collected near aquaculture facilities over the last two decades (O’Reilly *et al*. 2006; Porter *et al*.1998; Liu *et al*. 2017; Bradbury *et al*. 2020). The continued presence of European ancestry in North American aquaculture fish represents a significant elevation of both the potential threat and uncertainty associated with the impacts to already at-risk North American populations experiencing introgression from farm escapees (DFO 2016).

To date, the quantification of European ancestry in Atlantic salmon has been accomplished using small panels of microsatellite loci (King *et al*. 2001, O’Reilly *et al*. 2006) or large genomic panels (*e.g*., Liu *et al*. 2017; Bradbury *et al*. 2022). Accurate ancestry estimation requires extensive genome coverage but genotyping large numbers of individuals for thousands of markers can be cost prohibitive (Pucket 2017). In applied contexts, where the number of individuals may be large, a balance is therefore required to ensure that the genome is sufficiently sampled to allow for accurate admixture estimation, while keeping study costs reasonable. Studies characterizing the ability of different marker panels to accurately estimate admixture have repeatedly shown that larger panels, commonly comprised of hundreds or thousands of single nucleotide polymorphisms (SNPs), vastly outperform smaller panels of microsatellite markers (simple sequence repeats; SSRs) (Gärke *et al*. 2011; Camacho-Sanchez *et al*. 2019; Szatmári *et al*. 2021). The use of differing numbers of SSR and SNPs for differentiation of domestic chicken (*Gallus gallus*) breeds revealed that 70 SNP markers provided comparable performance to 29 SSR markers, while the use of 250 or more SNPs provided sufficient genomic coverage for accurate admixture estimation (Gärke *et al*. 2011). Similarly, repeated genetic clustering analyses for two amphibian species on the Iberian Peninsula showed that on data sets of similar sizes and spatial structures, tens of thousands of SNP markers outperformed panels of 18 and 14 microsatellites (Camacho-Sanchez *et al*. 2020). Ultimately, a comparison across marker types and a targeted screening tool for quantifying European introgression or individuals of European ancestry in Atlantic salmon is required and could provide the information necessary to mitigate some of the negative effects of aquaculture escapees on North American wild salmon populations.

Here we build on previous work identifying the presence of European introgression in farmed, escaped farmed, and wild Atlantic salmon throughout Atlantic Canada (*e.g*., Bradbury *et al*. 2022) and develop targeted genomic and machine learning tools to facilitate routine screening in support of conservation and management efforts. The goals of this study were to quantify the ability of panels of varying marker types (SSR and SNP), marker numbers, and panel designs to detect European introgression into North American farmed and wild Atlantic salmon, as well as to subsequently apply this information in the design of efficient tools for future *de novo* introgression detection. Specifically, we: 1) analyzed European admixture using three marker panels (7-SSR, 100-SSR, and 220K-SNP) on three different, but overlapping sets of thousands of Atlantic salmon; 2) used a common set of individuals to quantify the accuracy of European introgression detection by different panels relative to the complete genome-wide SNP marker panel (220K-SNP array); 3) isolated the effects of marker number, individual sample size, and the origins of individuals on admixture detection through down sampling and repeated admixture estimation; 4) designed, tested, and implemented a machine learning-based Python package with a Command Line Interface (CLI), SalmonEuAdmix, a diagnostic tool capable of accurately estimating European admixture proportions based solely on the genotype data of new samples for a set of 301-SNP markers, without the need for additional complete admixture analyses. The software SalmonEuAdmix is free and publicly available on GitHub (https://github.com/CNuge/SalmonEuAdmix)and the Python package index (https://pypi.org/project/SalmonEuAdmix/). The results demonstrate the power of targeted amplicons and machine learning algorithms to streamline ancestry estimation in support of the conservation of at-risk wildlife species.

## Materials and Methods

### Sample information & Genotyping

To compare the ability of different marker panels to detect European introgression into North American aquaculture and wild individuals, three data sets were utilized as the basis of the comparisons (Table 1). The first data set (220K-SNP) was a series of 7739 samples (Table 1) that were genotyped using a 220K bi-allelic SNP Affymetrix Axiom array developed for Atlantic salmon as described in Barson *et al*. (2015). Most of the samples utilized were from previously published sources (Lehnert *et al*. 2020; Bradbury *et al*. 2022), and all samples were subjected to the extraction, genotyping, and bioinformatics procedures described therein. The second data set (100-SSR) was a series of 3733 samples (Table 1) from a previously published source (Bradbury *et al*. 2018) that were genotyped using a panel of 100 microsatellite markers. This data set included wild and aquaculture fish from North America, but unlike the 220K SNP array data set it had European individuals exclusively derived from Norwegian aquaculture facilities (Table 1). The third data set (7-SSR) utilized was a series of 1516 individuals (Table 1) genotyped using a panel of seven microsatellite markers initially described in King *et al*. (2001). The samples genotyped for the 7-SSR panel were composed of wild and aquaculture samples from North America, as well as 269 triploid aquaculture individuals of European origin. Prior to genetic admixture analysis, the genotypes of the triploid individuals were down sampled. For each marker in each triploid individual, 2 of 3 alleles were randomly retained so as to create synthetic diploid samples suitable for use in subsequent admixture analysis. The three datasets had no overlap in genetic markers, but did have individual samples in common. A series of 370 individuals (211 North American aquaculture and 159 North American aquaculture escapees) were common to all three data sets and were used as a common test set for comparison of admixture detection across the different data sets.

**Table 1.** Origin of the Atlantic salmon samples genotyped using the different marker panels and utilized in the comparative admixture analyses. ^†^ The Unknown category were wild caught fish from New Brunswick, Canada of unknown wild or aquaculture origin. ^‡^ These were triploid samples that were genetically down sampled to create artificial diploids (2 of the 3 alleles were retained at random for each marker) for use in admixture analysis.

|  | North American |  |  |  |  | Icelandic | European |  |  |
| --- | --- | --- | --- | --- | --- | --- | --- | --- | --- |
| Data Category | Wild | Aquaculture | Aquaculture Escapee | Aquaculture Wild Mix | Unknown<sup>†</sup> | Aquaculture Iceland | Aquaculture Norway | Norway wild | Total |
| 220K-SNP panel | 5570 | 440 | 496 | 195 | 27 | 18 | 187 | 806 | 7739 |
| 100-SSR panel | 2733 | 201 | 296 |  | 44 |  | 187 + 272<sup>‡</sup> |  | 3733 |
| 7-SSR marker panel | 614 | 195 | 385 |  | 44 |  | 269<sup>‡</sup> |  | 1516 |

### Detection of European introgression through admixture analysis

For the 220K-SNP data set the data quality control (QC) filtering steps described in Bradbury *et al*. 2022 were replicated. We then conducted a principal component analysis (PCA) of the 220K-SNP marker genotypes using the program *pcadapt* (Luu *et al*. 2016) to quantify the population structure of the samples and ensure that the patterns described in Bradbury *et al*. (2022) were replicated in the present study. The program *Admixture (version 1.3.0*; Alexander *et al*. 2009) was then used with the parameter *k = 2* to calculate per individual admixture values; these results were visualized in R and admixture populations were retained for subsequent comparative analyses. For both the 100-SSR and 7-SSR panels the program *Structure (version 2.3.4;* Pritchard *et al*. 2000) was used to calculate admixture proportions for each individual (with the parameters: *k = 2, burn in = 50000 iterations, repetitions = 500000*). This change in admixture calculation method was required because unlike *Admixture*, *Structure* can accommodate microsatellite data with three or more alleles per locus, while yielding similar results (Alexander *et al*.2009).

### Comparison of admixture estimates across marker panels

The admixture proportion predictions made using the complete 220K-SNP marker panel (~186K markers post filtering) were assumed to be the most accurate measure of European introgression due to having the most comprehensive marker coverage and largest baseline sample sets of wild and farmed North American and European individuals. To assess the relative performance of the 220K-SNP and the two SSR marker panels, the per-individual estimated admixture proportion values (Q1 and Q2 estimates) for the set of 370 individuals common to all the data sets were considered. For both the SSR marker panels, a linear regression was performed with the European admixture proportion predicted by the complete 220K-SNP marker set used as the response variable and the European admixture proportion of the given SSR set used as the predictor. The regression coefficient (*r^2^*) was considered to be the measure of how well the complete ground truth admixture proportions were replicated by the predictor data set and the per individual predictions were visualized via scatterplots in R. Prediction accuracies were also examined from the perspective of a classification problem. Ground truth and predicted admixture values were converted to binary classifications, using a threshold of 0.1 to classify whether an individual was of pure North American origin (<0.1) or displayed significant European ancestry (≥0.1); these classification data were then used to generate confusion matrices and calculate prediction accuracy scores and error rates.

The use of three overlapping but non-equivalent sets of individuals for admixture prediction by the different marker panels provided a confound that prevented the regression coefficients and admixture proportions from being compared in a completely equivalent fashion, as the sample number and marker number did not vary independently. To better understand and quantify the effects of marker number and sample number independently, we conducted several additional admixture prediction analyses aimed at trying to isolate these variables. To provide evidence of the role of marker number and coverage on the detection of European introgression, random down sampling was conducted to produce smaller panels from the 220K-SNP data set. Random genome-wide subsets of 500, 400, 300, 200, and 100 SNPs were chosen from the complete set of SNPs that passed the filtering steps. For each of the samples, the admixture analysis was then repeated and the results for the 370 common test individuals were compared to the 220K-SNPmarker set predictions via linear regression to quantify admixture prediction accuracy. Finally, we isolated the effect of sample size by repeating the admixture proportion prediction analysis using the full set of markers from the 220K-SNP panel, but down sampling to produce a sample set with smaller numbers of individuals from the North American and European baseline populations. Two smaller sample sizes were extracted from the full data set and individuals were selected to as closely as possible mirror the compositions of the individuals genotyped with the 100-SSR and 7-SSR panels (Table 1). Both subsets included the 370 common test individuals to allow for direct comparison to the other analyses and the remaining subset were randomly selected from the complete set of available individuals on a within-category basis (categories listed in Table 1). The larger subset (mirroring the 100-SSR data set) was composed of 3485 individuals, including 2733 wild North American samples, 177 North American wild caught individuals of aquaculture or wild-aquaculture mixed origin, and 205 Norwegian aquaculture samples. The smaller 1441 sample set consisted of the 370 common individuals as well as 614 North American wild, 252 North American wild caught individuals of aquaculture or wild-aquaculture mixed origin, and 205 European aquaculture samples. The most significant change to these two down sampled data sets relative to the full 220K sample set was the complete exclusion of the 806 wild European samples. Admixture calculations and regression analyses comparing these per sample admixture predictions to the complete data set were then performed.

The interaction of reduced markers and individual sets was then examined through additional admixture analysis runs using the 3485 and 1441 individual sets and the following marker sets: the 500 random SNP panel and the 100 random SNP panel. These admixture analyses were conducted to see if the effects of marker number and sample number on admixture prediction accuracy were additive, and to better understand the influences of these variables on admixture prediction. The admixture predictions for the 370 common individuals were compared to the results from previous tests, to give an indication of the performance difference when the marker and individual numbers are both reduced.

### Design and testing of SNP marker panel

Following the comparative study of the marker panels and assessment of the relative importance of marker number and sample size, a sub-panel of SNPs was designed with the goal of producing a standardized set of markers for future per-individual admixture estimation with good genome coverage and strong lab-based performance metrics. The panel, of 301 SNP markers was selected from the complete set of 220K array SNPs based on several criteria: i) markers had to pass all the QC filtering steps in the 220K-SNP admixture analysis, ii) markers were selected so as to guarantee that all 29 chromosomes of the Atlantic salmon genome were represented, iii) markers were selected that had associated DNA sequences that analysis with PrimerServer (Zhu *et al*. 2017) predicted to have specific amplicon targets, iv) markers were selected that had high FST in comparison of North American and European ancestry individuals. The panel was subset from the complete data set using *Plink* (*version 1.90*) (Purcell *et al*. 2007) and used to conduct an additional admixture analysis. The results were compared to the admixture proportions predicted using the 220K-SNP panel via a linear regression.

Classification-based comparison of predictions to the 220K-SNP panel predictions was also conducted, with predicted admixture proportions converted to binary classifications using a threshold of 0.1 (pure North American origin <0.1, European ancestry introgression >=0.1).

### Machine learning model and Software design

After describing the ability of the various marker panels to detect European introgression, we aimed to design a software tool in the Python programming language to allow an end user to obtain an admixture prediction based on the 301-SNP panel without the need to re-run a complete admixture pipeline for each new set of samples, thereby increasing the feasibility of admixture detection for ongoing salmon conservation efforts. The software would reduce the data processing and computational overhead needed to estimate the European admixture proportion for a new sample or set of samples. To accomplish this, we trained and tested a series of supervised machine learning models to predict European admixture proportion (y) based on the SNP genotypes of a new individual for the markers in the selected panel (X).

To interface with the machine learning models, the genotype data for the complete set of 7636 individuals was numerically encoded in dosage format. Data processing code (https://github.com/CNuge/SalmonEuAdmix) was developed to read in a genotype file (in *Plink’s* PED format), impute missing genotypes with the mode genotype from the 220K-SNP data set, and numerically encode the genotypes (AA = 0, AB = 1, BB = 2, where A is the major allele in the baseline data and B is the minor allele). The set of 370 common individuals used in previous analyses were withheld to serve as a final validation set. Of the remaining individuals, 80% of the remaining individuals were randomly selected to form the training set for the machine learning models and 20% were withheld to serve as a test set spanning all the available data classes. The 370 common individuals assessed performance only on North American aquaculture and wild fish, while the test set additionally included individuals of complete European origin. To eliminate potential bias and ensure that the 370 individuals in the final validation data were completely withheld prior to final model assessment, an additional admixture run was conducted using the 301-SNP markers and the 370 validation individuals removed. The European admixture proportions obtained from this admixture run were used as the response variables (y) in model training.

Within Python, the three machine learning models: a random forest (RF), a support vector machine (SVM), and a deep neural network (DNN), were fit to the training data and used to make predictions on the withheld test and validation data. The RF model was implemented using the *sklearn.ensemble.RandomForestRegressor* function of the package *Scikit-learn* (Version 0.24.2, Pedregosa *et al*. 2011) using an *n_estimators* parameter of 1000 and defaults for all other parameters. The support vector machine (SVM) was implemented using the *sklearn.svm.SVR* function of *Scikit-learn* using a C value of 1.0, and an epsilon value of 0.2, and defaults for all other parameters (Version 0.24.2, Pedregosa *et al*. 2011). The DNN was a custom architecture designed using the package *Tensorflow* (Version 2.8.0, Abadi *et al*. 2016) that featured an input layer shape of 301 (matching the SNP panel size) three hidden layers of 1026, 342, and 114 densely connected neurons using the rectified linear activation (relu) function activation and 0.2 dropout frequency, and a single neuron output layer using a linear activation function. Training of the DNN used the Adam optimization algorithm, 20 training epochs, and mean absolute error as the loss metric. Code for the DNN model architecture can be found within the SalmonEuAdmix package (https://github.com/CNuge/SalmonEuAdmix/blob/master/SalmonEuAdmix/model.py).

The models were all trained with a 1 x 301 predictor tensor containing the dosage encoded genotypes, and the European admixture proportions obtained from admixture analysis using the 301-SNP panel set as the response variable. The response variables were scaled using a *StandardScaler* (Scikit-learn Version 0.24.2, Pedregosa *et al*. 2011) that was trained on the training data and applied to each of the train, test, and validation response variable sets. Predicted values were compared to the ground truth admixture proportions (Figure 1, Figure S1) obtained using the 220K-SNP data set. For each model, the root mean squared error was calculated and the predictions were saved to a tab separated output file. These data were then loaded into R where linear regressions were performed to compare the models’ predicted admixture proportions to the original values. Comparison of the results from the three different models was then used to select the optimally performing model. The final models were saved and the software package SalmonEuAdmix (https://github.com/CNuge/SalmonEuAdmix) was created to allow for efficient model reuse via a CLI.

**Figure 1.**
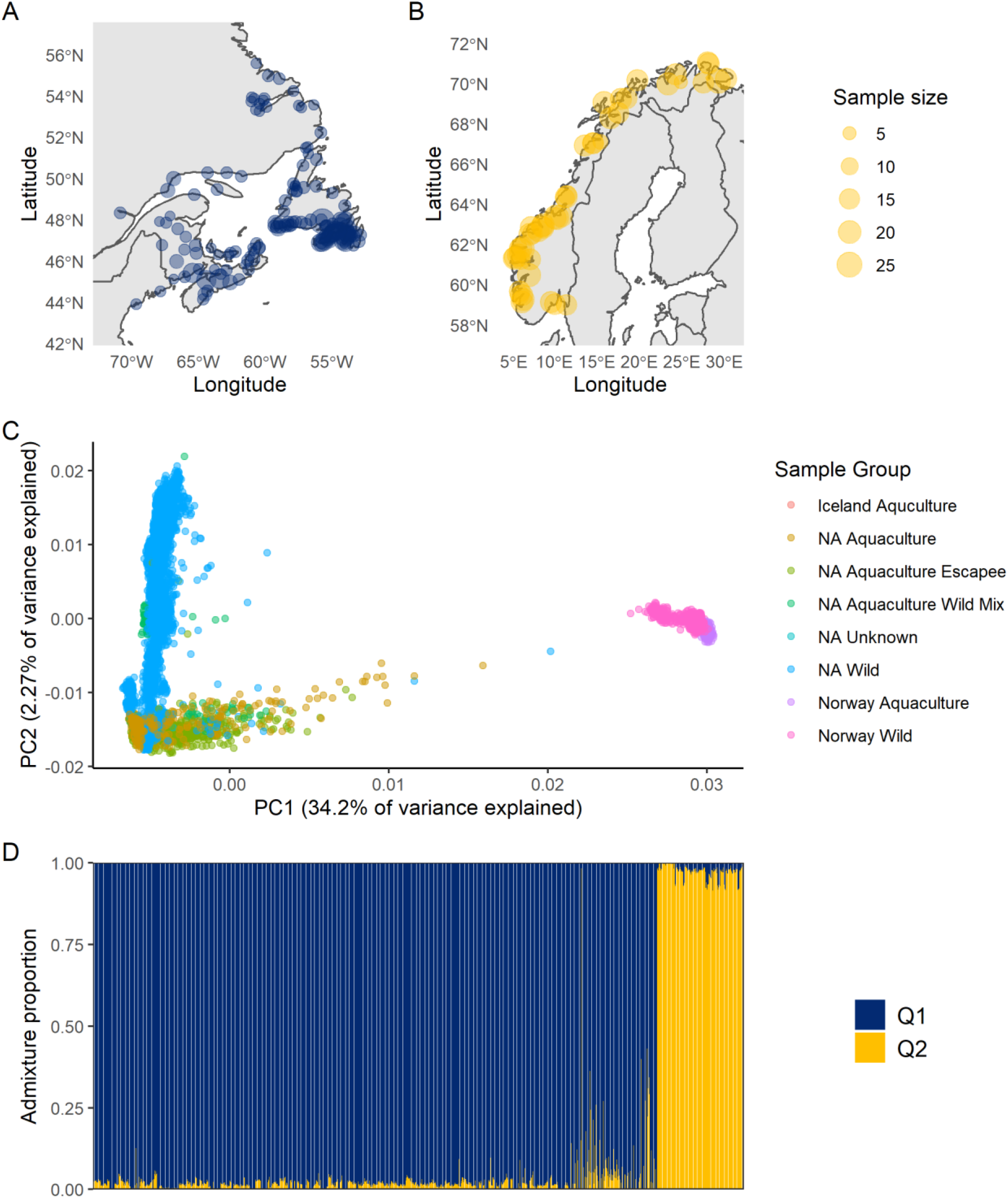
A) Map of the 148 sampling locations for the 5570 wild North American Atlantic salmon used in the study. B) Map of the 50 sampling locations for the 806 wild European Atlantic salmon used in the study. C) Scatter plot of Principal Components (PCs) of genetic variation for the 7636 Atlantic salmon genotyped using the 220K-SNP panel. The 186292 SNPs that passed quality control and filtering steps were the input for the PCA. The colour of the points indicates the category of origin for the samples (Table 1). D) Per-individual European admixture proportion estimates (Q2-values) based on admixture analysis of the 186292 SNPs passing quality control for the 7636 Atlantic salmon genotyped using the 220K-SNP panel. The samples are sorted by their data category of origin in the same left to right order as presented in Table 1.

## Results

### Detection of European introgression through admixture analysis

Following SNP and individual data filtering based on the criteria laid out in Bradbury *et al*. 2022, the 220K-SNP marker panel used for European admixture detection consisted of 186292 SNPs and 7636 individuals. Similar to the results reported in Bradbury *et al*. 2022 (with minor differences resulting from the increased sample size), the PCA revealed strong separation of samples of European and North American origin along the first axis of variation (PC1 = 34.2% variance explained), and evidence of individuals with mixed ancestry (Figure 1). The admixture analysis with the 220K-SNP panel separated North American wild fish from Norwegian fish of wild or aquaculture origin with high fidelity, while samples from the North American aquaculture and aquaculture escapee groups displayed evidence of European introgression (Figure 1).

For the 100-SSR marker panel, a total of 3646 individuals were successfully genotyped and passed all QC steps. The PCA showed the primary axis of variation was separating individuals of European and North American ancestry (PC1 = 6.6%, PC2 = 1.4% variance explained; Figure S2). The linear regression of the admixture proportions for the 370 commonly genotyped individuals revealed a significant, but weak concordance of predicted admixture proportions with the 220K-SNP panel predictions (r^2^=0.64 (Figure 2A). For the 7-SSR marker panel, 1438 individuals were genotyped and passed all QC steps. The PCA showed the primary axis of variation was separating individuals of European and North American ancestry (PC1 = 6.5%, PC2 = 1.9% variance explained; Figure S2). The linear regression of the 7-SSR admixture proportions for the 370 individuals showed lower concordance with the 220K-SNP panel admixture proportions predictions (r^2^=0.49). Inspection of the 7-SSR admixture proportion predictions for the 370 individuals showed a high number of individuals predicted to have less than 1% (242/370 = 66% of individuals) of European ancestry, while the 220K-SNP data set reported only 151 individuals with European admixture proportions less than 1% suggesting reduced ability to detect European admixture with the 7-SSR marker panel set. (Figure 2B).

**Figure 2.**
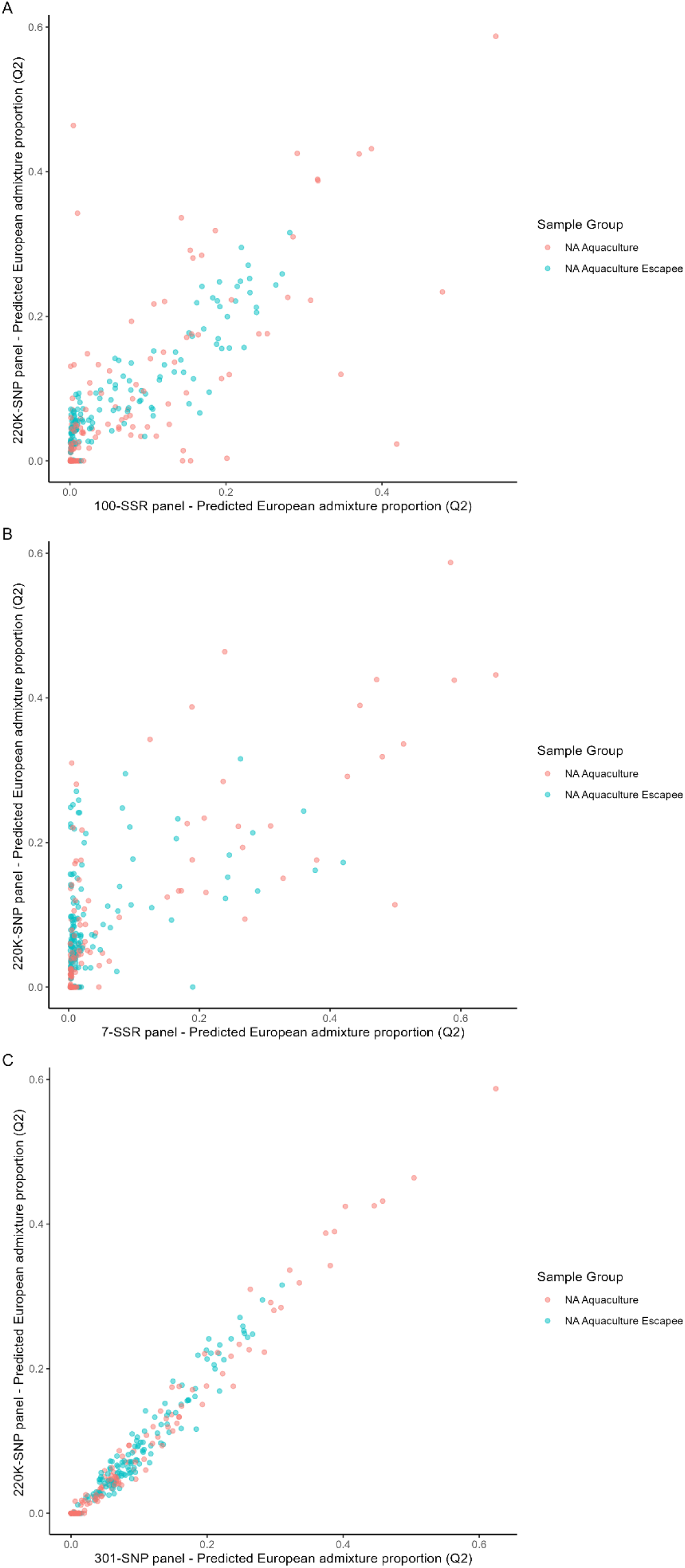
A) Scatter plot comparing the per-individual European admixture proportion predictions made by the 100-SSR SNP panel (x-axis) to the European admixture proportion predictions made using the 220K-SNP panel for the 370 individuals common to the two data sets. The colour of the points indicates the category of origin for the given sample. B) Scatter plot comparing the per-individual European admixture proportion predictions made by the 7-SSR SNP panel (x-axis) to the European admixture proportion predictions made using the 220K-SNP panel for the 370 individuals common to the two data sets. The colour of the points indicates the category of origin for the given sample. C) Scatter plot comparing the per-individual European admixture proportion predictions made by the 301-SNP panel (x-axis) to the European admixture proportion predictions made using the 220K-SNP panel for the 370 individuals common to the different marker panel data sets. The colour of the points indicates the category of origin for the given sample, Adjusted R-squared: 0.9754, p < 2.2e-16.

### Separating marker and sample effects

A series of additional admixture detection runs were conducted to isolate the effects of marker number and individual number on the characterization of European admixture. First, we isolated the effect of marker number by conducting random down sampling of SNPs while keeping the number of individuals constant (n=7636). Linear models were used to obtain the regression coefficients for each of the random marker subsamples (Table 2; Figure 3). The 500 random SNP marker panel performed better than either SSR panel, reproducing the 220K-SNP admixture predictions with an r^2^ of 0.97. The 400 and 300 random marker panels also had regression coefficients of greater than 0.95, suggesting that these marker sets had sufficient genome coverage to replicate the 220K-SNP admixture predictions with greater than 95% accuracy. The 200 random SNP panel displayed a larger performance decline relative to the larger random panels, with an r^2^ of 0.91 and the 100-SSR panel displayed lower performance still, with r^2^ of 0.83.

**Table 2.** Summary of regression results for the comparison of the predicted admixture proportions from different marker panels to the admixture predictions made using the 220K-SNP data set for the common set of 370 individuals.

| Analysis purpose | Panel used for admixture prediction | Number of markers | Number of individuals | r<sup>2</sup> when compared to 220K-SNP panel admixture proportions |
| --- | --- | --- | --- | --- |
| Panel comparison | 100-SSR | 100 | 3733 | 0.6432 |
|  | 7-SSR | 7 | 1516 | 0.4858 |
| Quantifying marker number effect | 500 random SNP | 500 | 7636 | 0.9684 |
|  | 400 random SNP | 400 |  | 0.9576 |
|  | 300 random SNP | 300 |  | 0.9507 |
|  | 200 random SNP | 200 |  | 0.9131 |
|  | 100 random SNP | 100 |  | 0.8292 |
| SNP sub-panel design | 301-SNP | 301 | 7636 | 0.9754 |
| Quantifying sample number effect | 220K-SNP – down sampled individuals | 186292 | 3485 | 0.9982 |
|  |  |  | 1441 | 0.9968 |
|  | 500 random SNP – down sampled individuals | 500 | 3485 | 0.969 |
|  |  |  | 1441 | 0.9696 |
|  | 100 random SNP – down sampled individuals | 100 | 3485 | 0.8159 |
|  |  |  | 1441 | 0.8424 |

**Figure 3.**
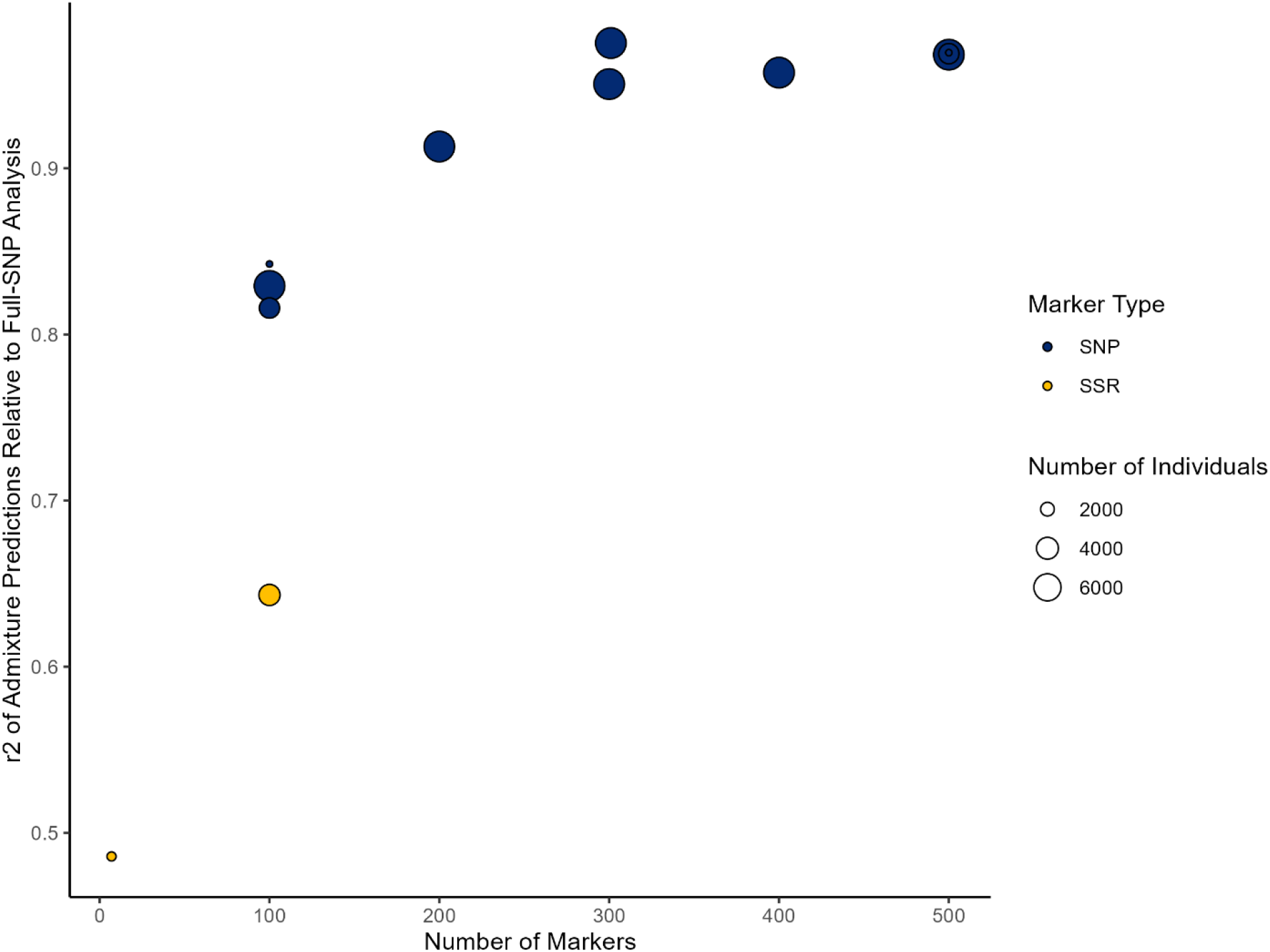
Scatter plot comparing the predicted admixture proportions from different marker types, marker numbers, and individual sample sizes to the admixture predictions made using the 220K-SNP data set for the common set of 370 individuals. Exact sample size, marker numbers, and r^2^ coefficients are presented in Table 2.

A second series of additional admixture analyses were run to isolate the effect of individual sample size on the characterization of European Admixture. For these tests, the composition of the number of individuals in the dataset was changed to resemble the number and type of individuals genotyped with the 100-SSR and 7-SSR panels (the data were down sampled to 3485 and 1441 individual sets respectively). Admixture analyses were run for these down sampled individual sets using: the 220K-SNP marker panel, the 500 random SNP panel, and the 100 random SNP panel. For each panel, when the number of individuals used in the admixture analysis was reduced there were no significant reductions observed in the correlation of the admixture prediction values, and those obtained using the 220K-SNP data set (Table 2; Figure 3). These results suggests that the number of markers had a larger impact on admixture detection than the number of individuals used in the admixture analysis.

### Testing of SNP marker panels

The PCA of the targeted 301-SNP panel produced genetic clustering patterns highly similar to the 220K-SNP panel, with strong separation of European and North American origin samples along the primary axis of variation (301-SNP: PC1 = 13.1%, PC2 = 5.2% variance explained; Figure S2). The admixture analysis was repeated for the down sampled 7636 individuals using the 301-SNP panel and linear regression comparing the per-individual predictions to the 220K-SNP per-individual admixture predictions showed that the 301-SNP panel outperformed the SSR panels and the 500 random SNP panels, with and r^2^ value of 0.98 (Table 2; Figure 2C).

### Assessment of panel classification accuracy

Classification-based comparison of the admixture predictions of the 301-SNP, 100-SSR, and 7-SSR panels to the 220K-SNP panel predictions was conducted using a binary prediction threshold of 0.1 (pure North American origin <0.1, European ancestry introgression ≥0.1). The 301-SNP panel had the lowest mis-classification rate of the three panels, with a 4.8% error rate (Table 3A). The 301-SNP panel displayed sensitivity to European admixture, with only 3 false negatives and 15 false positives. The 100-SSR panel had a mis-classification rate of 9% (Table 3B), so although the per individual admixture values may not as strongly correspond to the 220K-SNP panel predictions, the population level characterization of the number of fish with European ancestry is similar (with 15 false positives and 18 false negatives). For the 7-SSR panel there is a 13.2% mis-classification rate, that was directional in nature with 46 false negatives and only 3 false positives (Table 3C).

**Table 3.** Confusion matrices comparing the number of samples with predicted European admixture proportions greater than or less than 0.1 for: A) the 220K-SNP panel and the 301-SNP panel, B) the 220K-SNP panel and the 100-SSR panel, and C) the 220K-SNP panel and the 7-SSR panel.

|  |  | 301-SNP panel classification |  |
| --- | --- | --- | --- |
|  |  | Predicted low European ancestry (Q2 < 0.1) | Predicted high European ancestry (Q2 >= 0.1) |
| 220K-SNP panel classification | Predicted low European ancestry (Q2 < 0.1) | 272 | 15 |
|  | Predicted high European ancestry (Q2 >= 0.1) | 3 | 80 |

B)
|  |  | 100-SSR panel classification |  |
| --- | --- | --- | --- |
|  |  | Predicted low European ancestry (Q2 < 0.1) | Predicted high European ancestry (Q2 >= 0.1) |
| 220K-SNP panel classification | Predicted low European ancestry (Q2 < 0.1) | 272 | 15 |
|  | significant European ancestry (Q2 >= 0.1) | 18 | 65 |

C)
|  |  | 7-SSR panel classification |  |
| --- | --- | --- | --- |
|  |  | Predicted low European ancestry (Q2 < 0.1) | Predicted high European ancestry (Q2 >= 0.1) |
| 220K-SNP panel classification | Predicted low European ancestry (Q2 < 0.1) | 304 | 4 |
|  | Predicted high European ancestry (Q2 >= 0.1) | 47 | 38 |

### Machine learning model comparison

Prior to training of the machine learning models we removed potential bias by producing blind admixture values (withholding the 370 validation individuals at all stages and reconducting the 301-SNP admixture analyses) for use as response variables in machine learning model training. A linear regression demonstrated that the blind admixture proportions did not differ from the per-individual admixture proportions (r^2^ > 0.99, p < 2e-16).

Following model training (using the test set and blind admixture values), predictions were made on the test and validation sets. The root mean squared error (RMSE) of predictions for the 301-SNP panel models on the test set (n = 1454, 20% of individuals) were: 0.0417 for the DNN, 0.013 for the RF, and 0.035 for the SVM. For the 301-SNP panel model’s predictions on the validation data the RMSE were: 0.018 for the DNN, 0.039 for the RF, and 0.035 for the SVM. The per-individual admixture predictions produced by the three models were then compared to the ground truth admixture values obtained using the full set of SNP markers and individuals (Figure 4). For both the test and validation data sets, the DNN output admixture predictions that most closely resembled the ground truth predictions with regression coefficients (r^2^) of 0.99 and 0.95 for the test and validation data respectively. The SVR performance was similar for both data sets (test r^2^ = 0.99, validation r^2^ = 0.95), and the RF model had comparable performance to the other models on the test data (r^2^ = 0.99), but inferior performance on the validation data set (r^2^ = 0.81), suggesting the RF had either overfit to the training data or that it was less effective at characterizing intermediate admixture values that were more prevalent in the validation data. The strong test set scores for of all models are likely due to the similarity of the training and test individuals, which were subsets of the original full set of 7636 individuals and contained samples of similar origin (*i.e*. individuals from same wild sampling locations or individuals derived from the same aquaculture stock) and also due to the test set having individuals with less admixed genomes (full European or North American origin). The validation individuals were completely withheld in the machine learning process (not included in the additional admixture analysis used to create response values for model training) and there was a higher proportion of intermediate admixture individuals compared to the test set which had many individuals of pure North American or European origin, making these values a more robust assessment of model performance.

**Figure 4.**
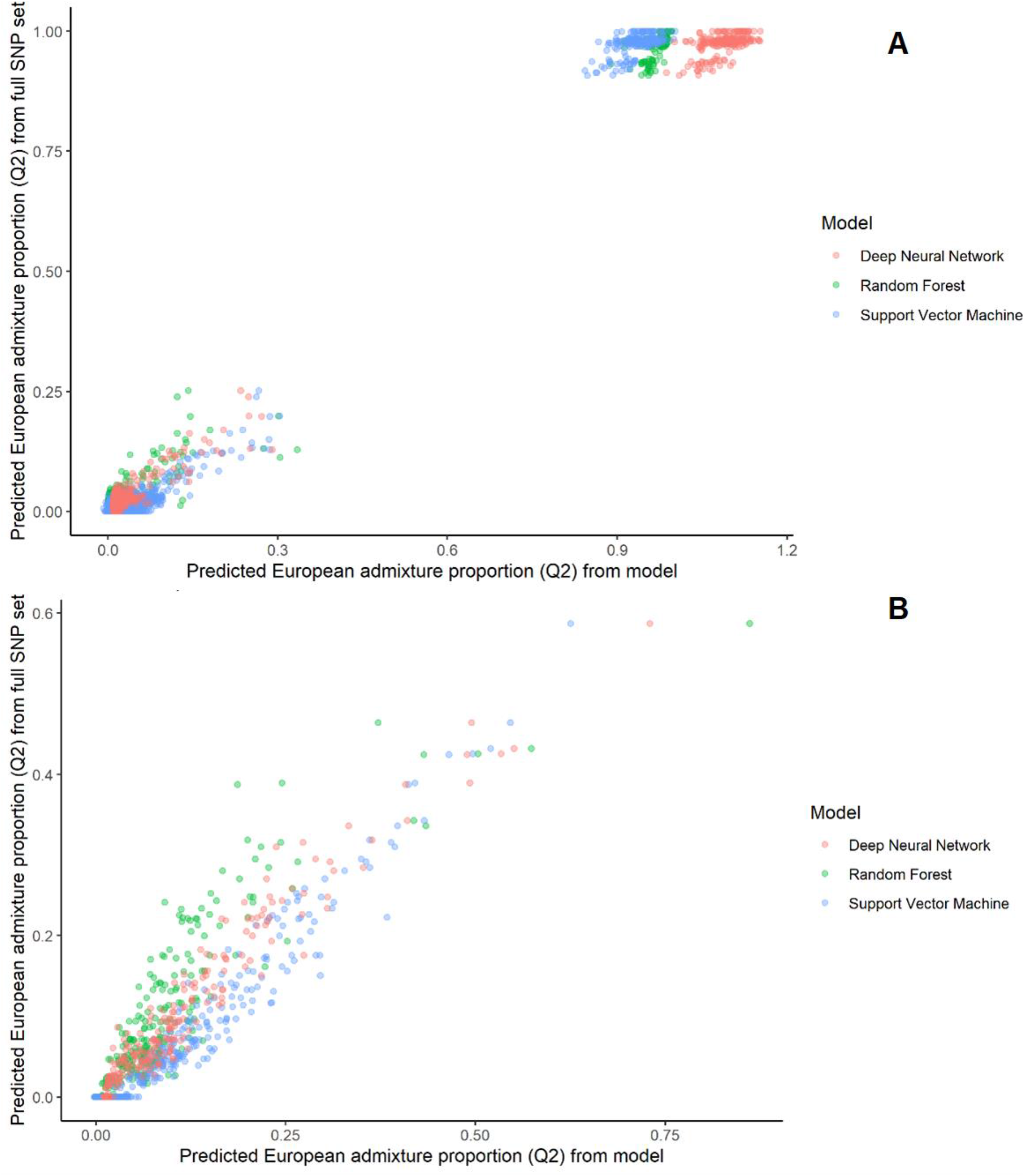
Scatter plots comparing the per-individual European admixture proportion predictions made by the three machine learning models (x-axis) to the original admixture proportion predictions made using the 301-SNP panel (y-axis) for: A) the 1454 randomly selected individuals in the test data set (r^2^ of regressions: Random Forest = 0.9973, SVM = 0.9948, DNN = 0.9980), and B) the validation set of 370 individuals common to the different marker panel data sets. (r^2^ of regressions: Random Forest = 0.8134, SVM = 0.9458, DNN = 0.9486).

Based on these results, the 301-SNP DNN model was selected for use in the SalmonEuAdmix package because of its ability to yield predictions that most closely resembled the European admixture proportions obtained through the complete admixture analysis for the previously unseen individuals. Due to the unconstrained nature of the DNN (*e.g*. predictions could be <0.0 or >1.0) there were individuals in the test set with predicted European ancestry proportions in excess of 1.0 (Figure 4). To account for this, a default, but optional heuristic was included in the SalmonEuAdmix package which constrained admixture predictions to a lower bound of 0.0 and an upper bound of 1.0.

## Discussion

Targeted SNP panels and admixture detection algorithms are becoming common place in conservation management activities revealing both population structure and hybridization (Camacho-Sanchez *et al*.2019; May *et al*. 2020; Stronen *et al*. 2022). In Atlantic Salmon, the identification of introgression of aquaculture salmon has become central to conservation efforts aimed at curbing salmon decline across the North Atlantic (*e.g*., Forseth *et al*. 2017; Bradbury *et al*. 2020) and genomic tools have been successfully applied to quantify hybridization and introgression (*e.g*., Karlsson *et al*. 2011; Pritchard *et al*. 2016; Wringe *et al*. 2019). Here we extended previous observations of aquaculture associated European introgression into North American salmon populations (O’Reilly *et al*. 2006; Bradbury *et al*. 2022) and develop targeted genomic and machine learning tools to mobilize European ancestry detection to inform conservation and management efforts. Our results suggest that accurate aquaculture associated European admixture estimation is possible with subsets of loci and accuracy is dependent more on genome coverage than number of baseline individuals considered. Iterative down sampling suggests that approximately 300 markers provided sufficient genomic coverage to closely replicate genome-wide admixture analysis in an efficient and cost-effective manner and that accuracy declined below this panel size. Combining this information with bioinformatics and lab-based metrics, we designed a panel of 301 SNPs, for use in future analyses aimed at characterizing European admixture proportions in North American populations. This panel, along with the deep neural network contained in the software package SalmonEuAdmix, allow for rapid and accurate *de novo* admixture proportion estimates to be made as part of future Atlantic salmon conservation and management efforts. The methods developed here serve as an example of how admixture data for at-risk wildlife species can be used in conjunction with machine learning algorithms to streamline ancestry estimation in support of conservation.

### Marker panel comparison

This work provides a comprehensive comparative study of the ability of different marker panels to detect European admixture within North American Atlantic salmon. The ability of the SNP array to accurately estimate individual ancestry was demonstrated through consistent performance across a range of marker panel sizes and baseline sample numbers. This is likely in part due to the high levels of differentiation between the North American and European lineages, which are estimated to have been isolated from one another for the past 600,000 years, with minimal secondary contact (Bourret *et al*. 2013; Moore *et al*. 2014; Rougemont & Bernatchez 2018; Lehnert *et al*. 2020; Bradbury *et al*. 2022). The inability to detect low levels of admixture was a limitation of the SSR panels (*i.e*., the 100-SSR and 7-SSR panels) as both of these SSR panels displayed reduced ancestry prediction accuracy (*i.e*. lower regression coefficients) compared to the 220K-SNP panel. These results for the 7-SSR panel are consistent with the hypothesis that the reduced performance of the SSR panels is mostly likely due to poor coverage of the Atlantic salmon genome. The Atlantic salmon genome has 27-29 chromosomes (Lien *et al*. 2016), so even if each of the 7-SSR panel’s markers were on separate chromosomes, any introgression on 22 of the 29 chromosomes (approximately 76% of the genome, or more depending on the size of the chromosomes containing the SSR markers) would not be in physical linkage with a panel marker and admixture in these regions would therefore go undetected. Scenarios with more European introgression, where recombination has occurred and smaller European ancestry tracts are present across numerous chromosomes, would go undetected by the 7-SSR panel unless by chance the admixture tracts span the SSR locations and contained a European ancestry tract. This same reasoning supports the major assumption we have made in the comparative study, which is that the 220K-SNP panel admixture predictions serve as a ‘ground truth’ to which other predictions are compared. With 186292 polymorphic SNP markers passing QC steps and being included in this panel, and the salmon genome being approximately 2.96 Gbp in size, the 220K-SNP panel provides genome wide coverage of approximately one SNP every 15.9 Kb of the Atlantic salmon genome, which is a level of genome-wide resolution sufficient to detect even very low levels of admixture (Lehnert *et al*. 2019; Bradbury *et al*. 2022).

Interestingly, the 100-SSR panel offered better genomic coverage than the 7-SSR panel, having specifically been designed to have representation of all chromosomes and therefore poor genomic coverage may not be the sole cause of its reduced admixture detection (Bradbury *et al*. 2018). An alternative hypothesis for the poorer performance of this panel relative to similarly sized SNP panels could be the accumulation of homoplastic (*e.g*. same repeat number) alleles within the North American and European lineages. Changes in microsatellite repeat number are a common mode of allelic evolution and have been shown to lead to microsatellite alleles of the same size with different evolutionary histories (Makova *et al*. 2000; Culver *et al*. 2001; Moodley *et al*. 2015). The estimated 600,000 YBP divergence time (Rougemont & Bernatchez 2018) of the two Atlantic salmon lineages would afford sufficient time for the accumulation of homoplastic microsatellite alleles and thereby contribute to the observed reduced admixture detection in comparison to the 100 locus SNP panel (see below).

The classification-based comparison of predictions further highlighted the differences in sensitivity to European admixture detection among the panels and demonstrated the potential impacts of these differences on classification-based screening of populations. Although the 7-SSR panel has previously been shown to have 100% correct continent of origin assignment (King *et al*. 2001), our work demonstrates that its capacity to detect European introgression is much more limited. The 7-SSR panel was shown to drastically under classify European introgression, which suggests that screening based on this panel would fail to detect European admixture in the majority of cases. Conversely, the 301-SNP panel possessed an error profile more suitable for applied conservation efforts aimed at screening for European admixture. The 301-SNP panel was sensitive to European admixture, detecting over 95% of true positives, while showing low levels of false positives as well. This is more suitable for screening in applied conservation efforts, where the costs of false negatives (overlooking true admixture and its associated negative effects) outweigh the costs of false positives (additional sampling or analytical efforts of non-admixed populations).

Admittedly, the direct comparison of panel results was limited to a subset of individuals (n = 370). Although these represented only a small fraction of the complete data sets, the admixture proportions of these individuals captured the level of ancestry variation in the total dataset and as such were well suited to assess the sensitivity of the different panels across a range admixture levels. For example, the 220K-SNP panel European admixture proportion predictions for these individuals ranged from 0.0 - 0.587 with 136 individuals having values in the range of 0.01 - 0.1 (*e.g*., 1% - 10% European Ancestry). These values reflect the range of admixture detected in broader analyses of aquaculture salmon and escapees (Bradbury *et al*. 2022) and also represent low admixture proportions that panels with poor genomic coverage would be more likely to fail to detect. If the common test set included more individuals with high (or complete) European ancestry, then the SSR panels admixture predictions would have likely more closely resembled the 220K-SNP panel predictions. Resolution of low to intermediate admixture proportions is of interest in applied conservation efforts, so the 370 individual test set used in this work is reflective of the context in which these findings will be applied and therefore likely very appropriate.

### Marker and sample number effects on admixture prediction

The iterative down sampling of SNPs showed an approximately linear decline, until a sharper drop in admixture prediction performance that was observed when only 200 markers were used; this is consistent with the hypothesis that at this point genomic coverage was sparse enough that larger admixture tracts went undetected. These results are similar to previous studies of admixture estimation using different numbers of markers, which have shown several hundred SNPs to provide sufficient genomic coverage for accurate estimation in a wide variety of species and contexts, while smaller panels (*e.g*. <100 markers) can have reduced admixture estimation ability in many situations (Vähä & Primmer 2006; Gärke *et al*.2011; Oliveira *et al*. 2015; Puckett & Eggert 2016; Fischer *et al*. 2017; Saint-Pé *et al*. 2019). The use of approximately 300 SNPs in subsequent custom panel design and predictive admixture model construction were therefore selected to strike a balance between genome coverage, admixture detection accuracy, and marker parsimony. The results of this study have shown only fractional performance declines for the 301-SNP panel relative to the 220K-SNP panel that was several orders of magnitude larger (when all other variables are held equal). Compared to genotyping individuals with the complete 220K Atlantic salmon SNP array (Barson *et al*. 2015), the 301 SNP genotypes required for admixture prediction with the 301-SNP panel can be obtained more economically and efficiently using targeted genotyping methods such as Genotyping-in-Thousands by sequencing (GT-seq) (Campbell *et al*. 2015).

The differences in the samples genotyped using the SSR and SNP marker panels complicated the interpretation of the results. Here, we attempted to isolate and quantify this effect through a comparative study of the admixture analyses and the use of down sampling to change the composition of individuals considered therein. In addition to the by-individual down sampled admixture runs that did not reveal significant effects of individual sample size on admixture predictions, comparing the difference in performance between the 100-SSR marker set and the 100 random SNP set (in terms of replication of the 220K-SNP admixture predictions on the 370 common individuals) indirectly gives an indication of the effect of the samples considered. The 100-SSR panel produced an r^2^ of 0.64, while the 100 random SNP panel produced an r^2^ of 0.83 (Table S1). This 0.182 difference in performance is unexpected given the information rich (*e.g*. multi-allelic) nature of microsatellite markers relative to bi-allelic SNPs and is contrary to previous work that has shown an opposing relationship of performance differences between similarly sized SNPs and SSRs sample sets utilized in admixture analyses (Gärke *et al*. 2011). As an alternative to the previously discussed microsatellite homoplasy hypothesis, the difference in performance may result from the bias introduced by the random SNPs being a subset of the 220K-SNP set used to obtain the ground truth admixture values and matching sets of individuals being used in these analyses. We attempted to quantify this bias through the down sampling of individuals to match the composition of the 100-SSR and 7-SSR admixture analyses, but this did not lead to any significant declined in the r^2^ of predictions relative to the 220K-SNP set. Conservatively, the 0.18 r^2^ difference between the 100 random SNP and 100-SSR marker sets may therefore be considered an estimate of the bias in favour of the SNP panel results, due to the SNP panels not being truly blind to the data in the 220K-SNP admixture predictions that constituted our ground truth values. Nonetheless, even with this bias taken into account (*e.g*. if we state that the hypothetical r^2^ of the 100-SSR is near or slightly higher than the 100 random SNP r^2^ of 0.8292), based on the other results of this study the 301-SNP panel would still likely far exceed the SSR panels’ admixture detection ability if the samples analyzed with the different marker panels were completely equivalent.

### SalmonEuAdmix and application of machine learning models

Machine learning models have recently been leveraged to infer genetic ancestry and to allow for the reconstruction of complex admixture histories in situations where traditionally employed methods can encounter limitations (Villanea & Schraiber 2019; Fortes-Lima *et al*. 2021; Bilschak *et al*. 2021). Our work represents a novel, alternative application of machine learning algorithms in ancestry estimation; instead of trying to better resolve admixture estimates, we trained supervised machine learning algorithms to replicate admixture proportion estimates which themselves were produced using an unsupervised learning algorithm (Pritchard *et al*. 2000; Tarca *et al*. 2007; Alexander *et al*. 2009). The predictive models learn the patterns relating genotypes to admixture proportions in the training data and make novel admixture estimates based solely on the genotypes of new individuals. This shifts the bulk of the analytical burden from the end user onto the algorithm designer, thereby transforming admixture estimation from a complex bioinformatics analysis into a simplified screening test, which is ideal for use in applied conservation efforts. This approach can be replicated within other species in order to take a robust set of admixture predictions and produce a customized diagnostic tool for rapid and simplified species-specific admixture estimation tool for use in applied conservation efforts (Oliveira *et al*. 2015; Bilschak *et al*. 2021; Stronen *et al*. 2022).

It is important to remember that this supervised learning approach to admixture estimation is meant to complement, not replace, traditional unsupervised admixture estimation methods. As evidenced by our assessment of panel classification accuracy, supervised models (such as the DNN used in SalmonEuAdmix) can be developed that are sensitive to the presence of admixture, allowing for the detection of cases of interest within applied contexts. However, the fine scale admixture proportions are inferior to a complete admixture analysis run using a maximal amount of available genetic markers. Within the intended application as an admixture screening tool, SalmonEuAdmix is likely to be robust, being based on genetic data from thousands of Atlantic salmon that display a spectrum of admixture proportions. The ability of SalmonEuAdmix’s models to predict admixture of previously untested populations is uncertain and may vary depending on the details of the population in question; however, we expect it to be effective for sample from novel locations in Atlantic Canada given the wide-ranging set of wild North American samples used in this study and the significant proportion of genomic variation explained by North American and European divergence. Despite potential limitation of model generalizability, the DNNs of SalmonEuAdmix are likely to outperform admixture analyses based on the 7-SSR or 100-SSR marker panels, as the 301-SNP panel provides greater genomic coverage and is comprised of bi-allelic SNPs (providing a defined parameter space for variation, whereas SSR markers may be found in novel variants within new populations). As more genotyped Atlantic salmon samples are made available, we will monitor SalmonEuAdmix’s performance in a growing number of contexts through the comparison of model predictions to additional, complete admixture re-analyses. Should areas of underperformance be identified, we will update the underlying model of SalmonEuAdmix and document changes in order to ensure the package provides accurate European admixture proportion predictions in the widest possible set of populations.

### Conclusion

The use of aquaculture salmon with European ancestry in North America presents a continued threat to declining North American Atlantic populations (Glover *et al*. 2017; Wringe *et al*. 2018; Bradbury *et al*. 2020, 2022). Extending previous studies which designed marker panels for aquaculture introgression (King *et al*. 2001; Bradbury *et al*. 2018; Bradbury *et al*. 2022), here our results present a comparison of different marker panel’s ability to detect aquaculture associated European introgression and demonstrated the greater accuracy and resolution of large SNP panels compared to commonly employed microsatellite-based methods. With the aim of producing the genomic and analytical tools necessary for efficient European admixture detection in future applied conservation efforts, we quantified accuracy differences between SNP panels of various sizes and used this information to inform the design of an optimized SNP panel, comprised of 301 markers, that provided highly similar admixture estimates to the 220K-SNP panel using a more parsimonious data set. To further aid the application of these panels in Atlantic salmon conservation and management efforts we developed the Python package SalmonEuAdmix (https://github.com/CNuge/SalmonEuAdmix), which uses the panels and a corresponding deep neural network to generate accurate estimates of European admixture proportions without the need for complete admixture analysis pipelines. The panels and software we have designed and tested will aid in Atlantic salmon conservation by providing the resources necessary to screen wild and aquaculture populations for evidence of European admixture and thereby allow evidence-based management decisions to mitigate negative impacts on wild populations throughout North America. The results also demonstrate how machine learning algorithms can streamline ancestry estimation to support applied conservation efforts; these techniques can be applied to other species at risk, allowing existing genetic information to be used to train models that facilitate rapid admixture estimates to inform conservation efforts.

## Acknowledgements

The authors would like to thank staff of Fisheries and Oceans Newfoundland and Labrador Salmonids Section for assistance with sample collection. The acquisition of aquaculture salmon baseline samples was facilitated by G. Perry, C. Hendry, DFO Aquaculture Section Newfoundland Region, and by industry partners Cooke Aquaculture, Grieg NL Seafarms Ltd, and Northern Harvest Sea Farms. We also thank CIGENE and RPC for SNP and microsatellite genotyping, and Ian Paterson and Beth Watson at the Marine Gene Probe Lab, Dalhousie University assisted with laboratory analysis. Funding was provided through the Program for Aquaculture Regulatory Research of Fisheries and Oceans Canada, the Genomics Research and Development Initiative of Canada.

## Data Accessibility Statement

Software described are free and publicly available (https://github.com/CNuge/SalmonEuAdmix). Data used in these analyses were generated in previous studies (Bradbury *et al*. 2018; Lehnert *et al*. 2019; Bradbury *et al*. 2022).

## Author Contributions

The study was designed by IRB, TK, CMN. Analysis design was done by CMN, TK, MKB, BLL, SJL, IRB. Data processing, statistical analyses, and data visualizations were conducted by CMN, TK, and SJL. Initial manuscript preparation was done by CMN. Python package design and programming was done by CMN. Testing of the Python package was done by CMN, MH, MKB, BLL, SVB. All authors contributed to the revisions of the manuscript.

## SUPPLEMENTARY INFORMATION

**Supplementary File 1** (‘S1-Supplementary_Tables_and_Figures.docx’) Supplementary tables and figures for the manuscript.

