## Supplementary material for "Genomic and machine learning-based screening of aquaculture associated introgression into at-risk wild North American Atlantic salmon (*Salmo salar*) populations": S1-Supplementary_Tables_and_Figures

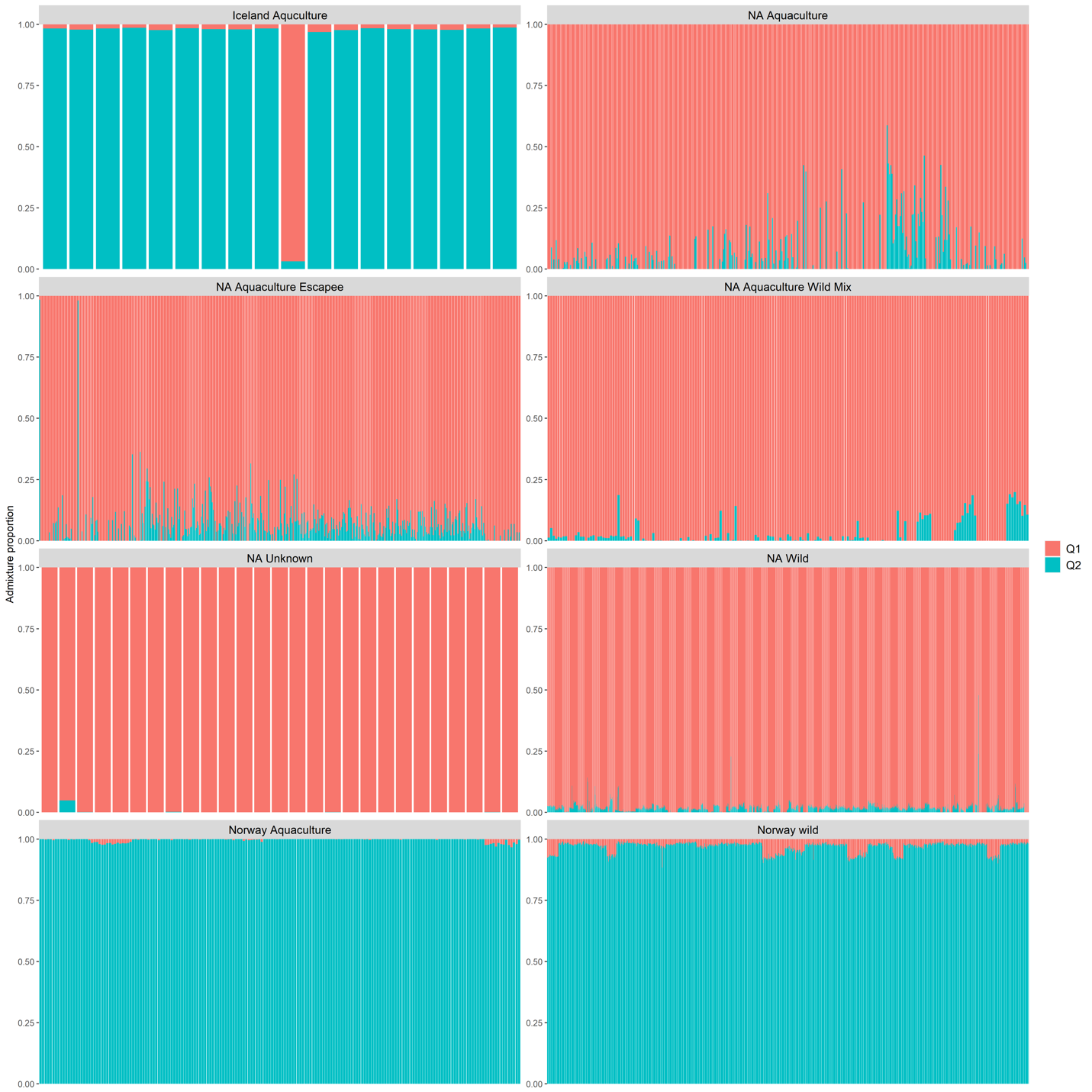

**Figure S1.** Per-individual European admixture proportion estimates (Q2-values) based on admixture analysis of the 220K-SNP panel (186292 SNPs post filtering) for the 7636 Atlantic salmon genotyped using the 220K SNP panel. The samples are separated by their category of origin (Table 1).

A

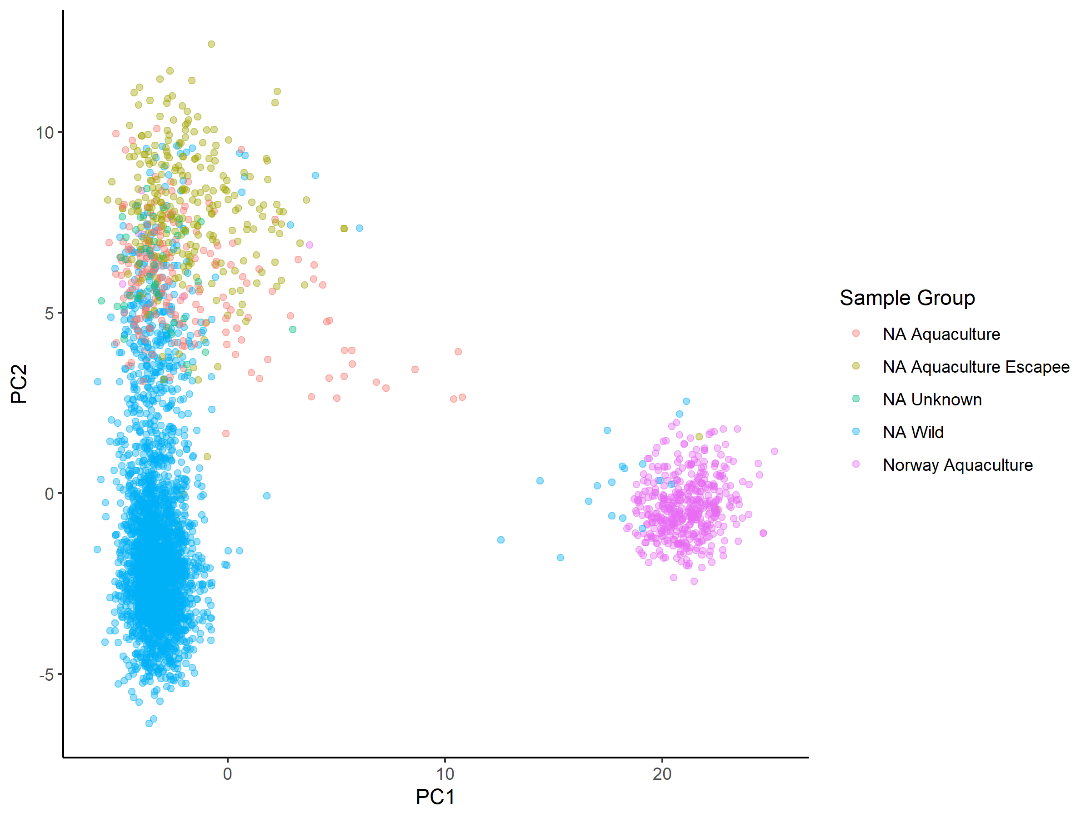

B

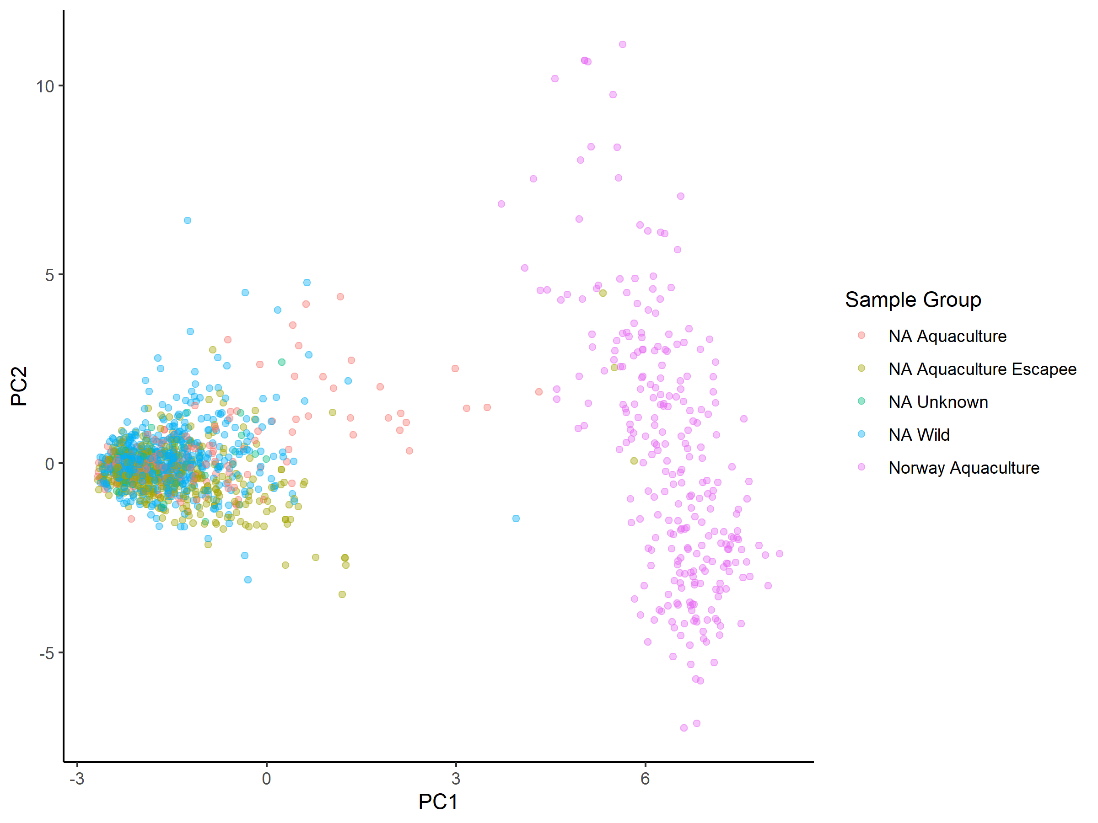

C)

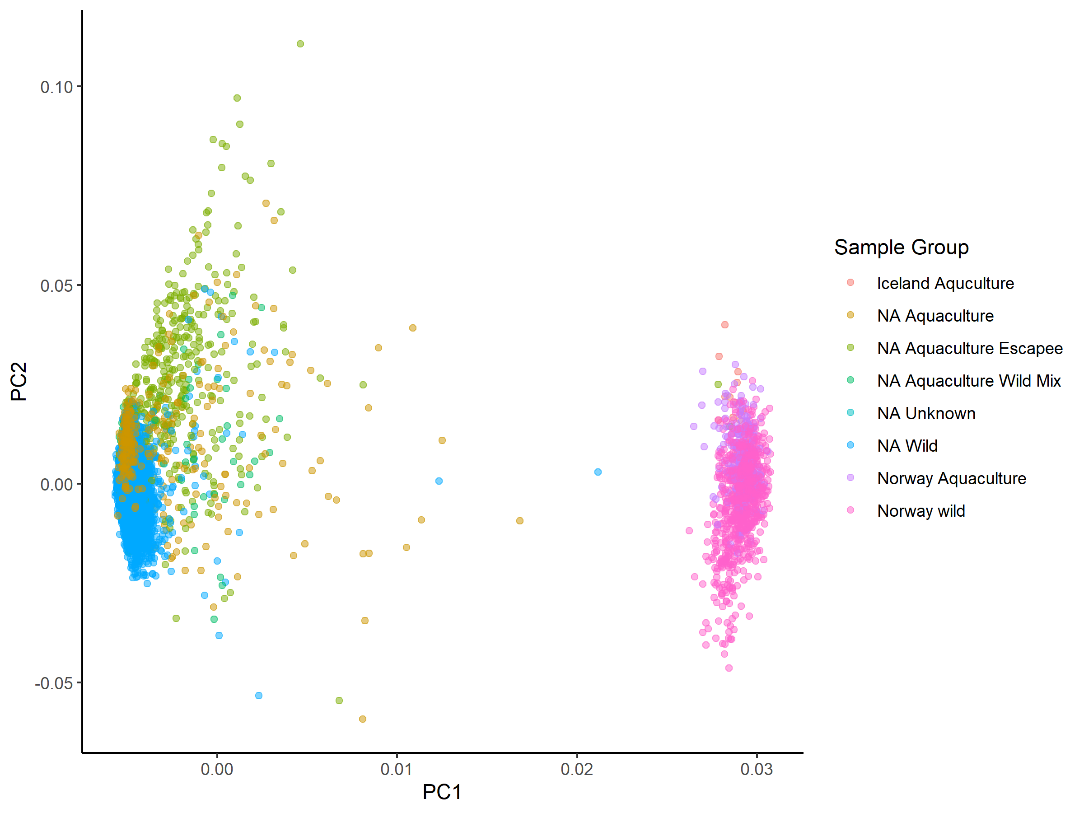

**Figure** **S2.** A) Scatter plot of PCs of genetic variation for the 3646 Atlantic salmon genotyped using the 100-SSR panel (PC1 = 6.6%, PC2 = 1.4% variance explained). The 100 SSR were dosage encoded and used as the input for the PCA. The colour of the points indicates the category of origin for the samples (Table 1). B) Scatter plot of PCs of genetic variation for the 1516 Atlantic salmon genotyped using the 7-SSR panel (PC1 = 6.5%, PC2 = 1.9% variance explained). The 100 SSR were dosage encoded and used as the input for the PCA. The colour of the points indicates the category of origin for the samples (Table 1). C) Scatter plot of Principal Components (PCs) of a PCA of genetic variation for the 301-SNP panel (PC1 = 13.1%, PC2 = 5.2% variance explained). The points for the 7636 samples are coloured by their category of origin (Table 1).

**Table S1.** Summary of regression results for the comparison of the predicted admixture proportions from different marker panels to the admixture predictions made by the 220K-SNP panel, with the linear regressions restricted to individuals with 220K-SNP admixture predictions < 0.1 and >= 0.1. ^†^ Indicates a linear regression where the regression coefficient had an associated p value greater than 0.05

| Admixture prediction source | Number of markers | Number of individuals | r^2^ for complete admixture proportions | r^2^ for just individuals with < 0.1 220K-SNP admix predictions  (n = 287) | r^2^ for just individuals with >= 0.1 220K-SNP admix predictions (n = 83) |
| --- | --- | --- | --- | --- | --- |
| 100-SSR panel | 100 | 3733 | 0.6432 | 0.17^†^ | 0.32 |
| 7-SSR panel | 7 | 1516 | 0.4858 | 0.086 | 0.2771 |
| 500 random SNP subpanel | 500 | 7636 | 0.9684 | 0.9273 | 0.8452 |
| 400 random SNP subpanel | 400 | 7636 | 0.9576 | 0.8961 | 0.8127 |
| 300 random SNP subpanel | 300 | 7636 | 0.9507 | 0.8970 | 0.7425 |
| 200 random SNP subpanel | 200 | 7636 | 0.9131 | 0.8247 | 0.6575 |
| 100 random SNP subpanel | 100 | 7636 | 0.8292 | 0.6684 | 0.4921 |
